# The effect of cathodal tDCS on fear extinction: a cross-measures study

**DOI:** 10.1101/561803

**Authors:** Ana Ganho-Ávila, Óscar F. Gonçalves, Raquel Guiomar, Paulo Sérgio Boggio, Manish Kumar Asthana, Angelos-Miltiadis Krypotos, Jorge Almeida

## Abstract

Extinction-based procedures are often used to inhibit maladaptive fear responses. However, because extinction procedures show efficacy limitations, transcranial direct current stimulation (tDCS) has been suggested as a promising add-on enhancer. In this study, we tested the effect of cathodal tDCS over extinction, to unveil the processes at play that boost the effectiveness of extinction procedures and its translational potential to the treatment of anxiety disorders.

We implemented a fear conditioning procedure whereby 41 healthy women (mean age = 20.51 ± 5.0) were assigned to either cathodal tDCS (n=27) or sham tDCS (n=16). Fear responses were measured with self-reports, autonomic responses, and implicit avoidance tendencies.

Cathodal tDCS shows no statistically significant effect in extinction, according to self-reports, and seems to even negatively affect fear conditioned skin conductance responses. However, implicit avoidance tendencies, assessed one-to-three months after the tDCS session and extinction, reveal a group difference in the avoidance tendencies towards the neutral stimuli (*F* (1, 41) = 12.04, *p* = .001, *ηp^2^* = .227), with the active cathodal tDCS groups showing a positive bias but not the sham group. This suggests a decreased generalization effect in the tDCS group with a moderate effect size. That is, cathodal tDCS may have enhanced long-term distinctiveness between threatening cues and perceptively similar neutral cues through a disambiguation process of the value of the neutral stimuli – a therapeutic target in anxiety disorders. Future studies should confirm these results and extend the study of cathodal tDCS effect on short term avoidance tendencies.

## Introduction

According to the classical fear conditioning model of anxiety disorders, anxiety and fear related responses result from associative learning processes, whereby threatening experiences are associated with originally neutral stimuli that subsequently gain anxiety-inducing properties (fear response acquisition). When individuals are subsequently exposed to these anxiety-inducing stimuli in safe conditions, a new learning occurs, the association between the stimuli and a potential threat is weakened, and the fear response is inhibited or eliminated (fear response extinction).

Current, exposure-based treatments for anxiety disorders are still limited in terms of their clinical efficacy with frequent recovery of symptoms (1,2). In this study, we used non-invasive transcranial direct current stimulation (tDCS) as an add-on intervention to enhance fear-extinction efficacy and observe its impact in three dependent measures that match three distinctive components of the fear response (3): subjective experience, autonomic responses, and implicit avoidance tendencies.

One promising way by which fear extinction procedures can be enhanced is through the use of non-invasive brain stimulation techniques. In particular, it has been shown that fear experiences can be modulated by tDCS (4,5). The modulatory assumptions are that tDCS 1) induces cortical excitability and neuroplasticity, modulating long-term potentiation (LTP) and long-term depression (LTD) mechanisms (6,7); 2) its effects are polarity specific (8,9) in that anodal stimulation is excitatory, whereas cathodal stimulation is inhibitory when stimulating motor or parietal regions but inconsistent in other brain regions (10); 3) it interferes with the cortical and subcortical regions involved in fear learning networks and their connectivity patterns (11,12); and 4) its effects may persist over time (13,14).

The modulatory effect of tDCS on the fear neural network, led researchers to consider this technique a potential booster to fear extinction procedures. The use of tDCS in extinction is based upon previous literature about the regions that participate in the fear-processing network. The vmPFC participates in fear learning by downregulating the amygdala activity during fear processing (15) and during fear extinction (16,17). Due to its regulatory effects, the vmPFC has been elected as a target region to modulate fear responses. However, tDCS currents cannot directly access this region, directing researchers to rely on the connectivity of the fear network and to identify other more accessible cortical regions such as the prefrontal cortex (PFC; 18,19), the DLPFC (4,19), or the supra orbital cortex (SOC; 5).

Recent studies observing anodal stimulation of the rDLPFC have led to conflicting results. Whereas Abend et al. (20) found anodal tDCS to have no impact in extinction, others found anodal tDCS to enhance both early (18) and late extinction (19). Moreover, both van’t Wout et al. (19) and Abend et al. (20) studies found a negative adverse effect of anodal tDCS as it seems to induce generalization of the fear response towards the neutral stimuli.

Similarly, albeit in laboratory cathodal stimulation of the rDLPFC of healthy subjects, showed no effect in fear extinction (21), two successful case reports found cathodal tDCS therapeutic effects (22,23), where we can see that cathodal stimulation may be a promising tool in alleviating anxiety symptoms. One possible explanation for the dissonant results between experimental studies and case reports comes from the different mechanisms that may be at play, whether extinction occurs inside or outside the reconsolidation time window.

Reconsolidation is thought to be the set of processes that occur after memory recall and during which a memory trace is destabilized. According to the reconsolidation theory, when its mechanisms are successfully triggered during its time-window the memory trace is labile and thought to be susceptible to be strengthened, updated or eliminated (24). In the case of the elimination of fear conditioned responses, the PFC structures participate in classical extinction out of the reconsolidation window, but are not necessary during reconsolidation (25). Previous literature has even suggested that the participation of the PFC during fear extinction when occurring within the reconsolidation time-window blocks the mechanisms involved in the long-lasting update of the conditioned fear response (25,26). Also, Abend et al. (20) suggested that cathodal stimulation may have a long-term depression effect over the medial PFC during the reconsolidation window, potentially eading to enhanced extinction. However, when Mungee et al. (21) used cathodal tDCS after fear recall (aimed at triggering reconsolidation), no impact was found on Skin Conductance Responses (SCRs). Of note, in their study, participants went through cathodal tDCS immediately after recall (a single presentation of the CS+), and the authors did not employ a fear extinction procedure, during which new information is thought to support the update of the fear memory trace, and to putatively lead to the persistent elimination of the conditioned fear response (27).

SCRs are slow autonomic responses to novel or emotionally salient stimuli, and the fear conditioned stimuli (CS+) typically elicits increased SCRs, compared to neutral conditioned stimuli (CS−). To test the tDCS effects on fear conditioning, most experimental studies use SCRs as indirect psychophysiological measures of conditioned fear (e.g. 4,5). For instance, Mungee et al. (5) observed that 24h after fear acquisition and after a CS reminder, anodal tDCS stimulation to the right PFC and simultaneous cathodal stimulation to the left SOC enhanced fear conditioned responses to the CS+, according to SCRs. In contrast, Asthana et al. (4) found that anodal tDCS delivered to the lDLPFC during fear acquisition consolidation period, had no impact on fear extinction 24 hours later, whereas cathodal tDCS had, leading to a decreased fear according SCRs.

Despite its extensive use, measuring fear responses relying solely on SCRs has its drawbacks. One of which is that this measure is sensitive to the repeated presentation of stimuli, resulting in a progressive decrease in SCRs amplitude (26,28). This habituation to the CSs is a phenomenon that compromises the length of the experimental procedures, the value of the signal across repeated sessions, and the usefulness of the results (29).

Notwithstanding, previous literature is relatively silent concerning the impact of tDCS in conditioned fear responses other than SCRs. Few exceptions look at gaze fixation times, associated with vigilance to fearful faces and found decreased fixation times subsequent to cathodal tDCS stimulation to the DLPFC (30). Neuroimaging studies showed that anodal tDCS over the DLPFC modulates vigilance to threatening stimuli (15), by means of a decreased amygdala activity. Nonetheless, the field has overlooked the implicit behavior component of the fear response when examining the efficacy of tDCS on extinction (31,32).

Implicit avoidance tendencies are argued to be automatic, involuntary, unconscious, goal-independent, and fast (33). Similarly to other components of fear, implicit avoidance can be acquired by associative learning and employed such that a threating forthcoming event is prevented (e.g. climbing a chair at the sight of a domestic spider;,3,34). Furthermore, when the fear of anxiety-inducing cues is generalized to stimuli that are perceptively similar to the original conditioned stimuli, individuals end up avoiding not only the conditioned stimuli but also a vast number of somehow related cues but otherwise harmless objects or situations (35).

Consequently, the presence of pervasive avoidance tendencies prevents individuals from experiencing the context as safe, hampering the success of fear extinction (36,37) by impairing new learnings, and prompting to fear recovery.

One hypothesis derived from the abovementioned studies is that by means of a reduced participation of the PFC in emotional regulation, cathodal tDCS may allow for a long-lasting fear memory update by disrupting reconsolidation processes (38,39). If this is true, then cathodal stimulation to the rDLPFC in tandem with a typical fear extinction procedure may lead to the extinction of autonomic fear responses, along with other resilient components of fear, such as avoidance tendencies. Furthermore, in parallel to the elimination of the fear response components, cathodal tDCS may result in the mitigation of the generalization effect towards perceptive or semantic similar stimuli (as unexpectedly happened in Abend et al. and Van’t Wout et al studies;,19,20).

In this study, we used a 3-day fear conditioning procedure to test whether cathodal tDCS over the rDLPFC enhances the efficacy of fear extinction procedures as assessed by three components of the fear response: the autonomic response (SCRs), the subjective experience (self-reports) and the implicit avoidance tendencies measured by reaction times (RTs; 40; cf. Fig 1).

**Fig 1.**
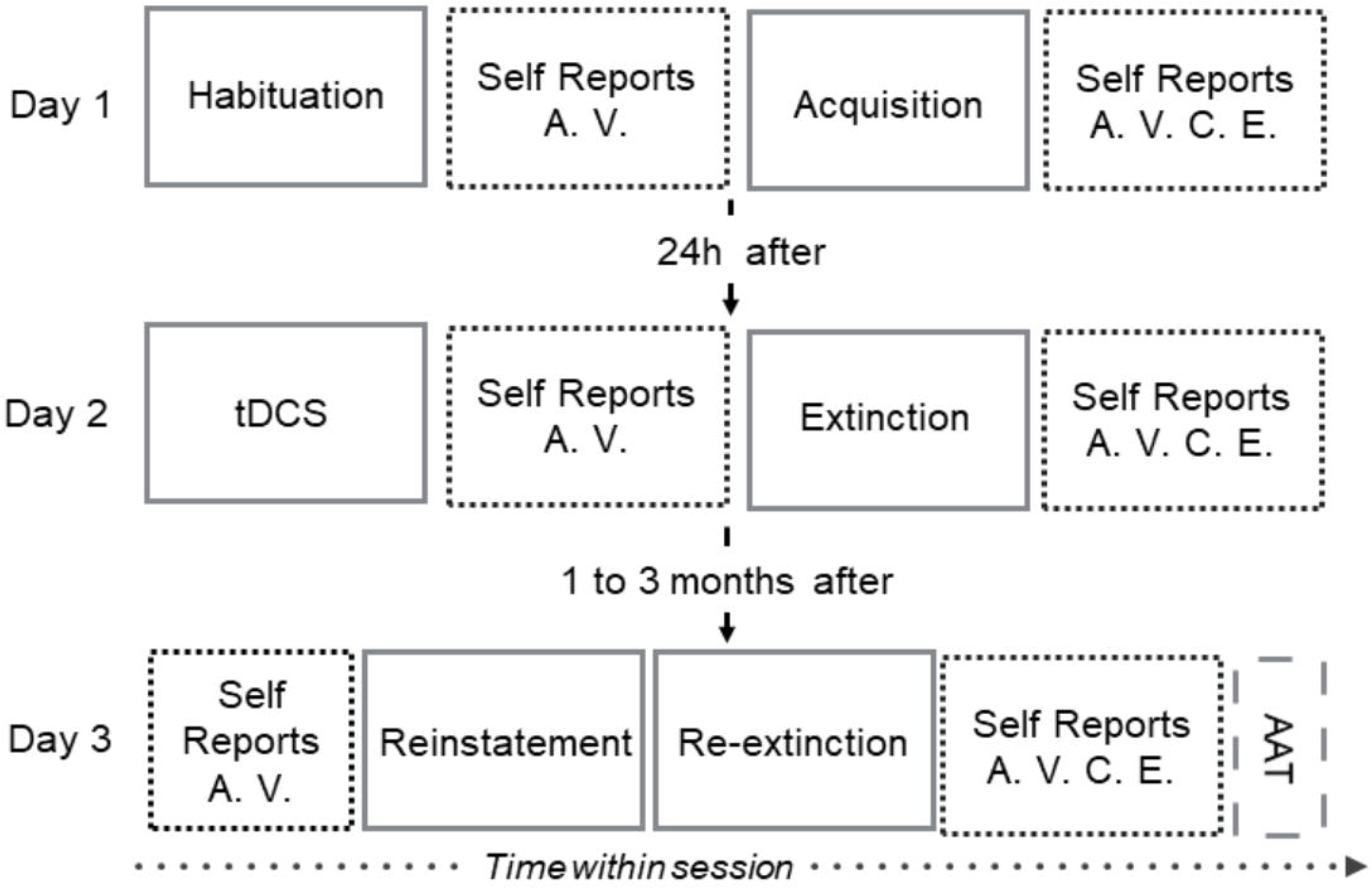
Experimental design. In day 1, participants went through habituation to stimuli (7 CS+; 7 CS−) and fear acquisition session (16 CS+, 16 CS−; 12 US). In day 2, participants were randomly assigned to the cathodal tDCS group or the sham tDCS group. The 20min tDCS session was followed by extinction training (16 CS+; 16CS−). In day 3, participants recovered the conditioned fear with a reinstatement procedure (4 US), followed by re-extinction (16 CS+; 16CS−) and the AAT (8 practice trials and 80 experimental trials. tDCS: Transcranial Direct Current Stimulation; AAT: Approach-avoidance Task; CS+: conditioned stimuli; CS−: unreinforced or control stimuli; US: Unconditioned stimuli. Self-reports: A – arousal; V – valence; C – contingency; E – expectancy. (Single column image, grey scales)

## Methods

Forty-eight women (mean age = 20.51 years, SD = 5.00) participated in the study, were informed about the procedure, and gave their written informed consent. The study, including its experimental protocol, was conducted according to the Declaration of Helsinki and approved by the Ethical Committee of the Faculty of Psychology and Educational Sciences, of the University of Coimbra (Ref. DIR352/2014). We defined the following exclusion criteria: < 18 years of age, history of psychiatric disorder and/or current psychoactive medication, screened by an experienced clinical psychologist using a short DSM-4-based clinical interview (41); pregnancy; caffeine and/or alcohol intake 24h before sessions; having had any physical exercise or meal 2h before the start of each session (42); auditory or visual (non-corrected) deficit; and contraindications to the use of tDCS (43). Additionally, participants had to show contingency awareness, as measured by self-reported contingency ratings (CS+/US ≥ 50% and CS−/US < 50%; see “Other self-report measures” section below).

### Stimuli

We used three types of stimuli– the unconditioned stimulus (US), the conditioned stimulus (CS+), and the neutral stimulus (CS−; 4, 16, 25). The CSs were blue or yellow 12×12cm squares (counterbalanced across participants), presented against a white background for 12s. The US was a women’s scream (item 277 from the International Affective Digitized Sound System, 4,44) delivered through noise cancelling headphones.

We pseudo-randomized the CS+ and the CS− trials within each session of each day, such that no more than two consecutive presentations of the same category of stimuli were allowed. For stimuli presentation, we used E-Prime (2.0.10.353 Standard SP1, Psychology Software Tools, Pittsburgh, PA), connected to a DELLP2012H monitor.

### Fear conditioning procedure

The experiment consisted of a partial-reinforcement auditory fear conditioning procedure, divided in two consecutive days (day 1–baseline psychological assessment questionnaires, fear habituation and fear acquisition; day 2–tDCS session and fear extinction) plus a follow-up session (day3–fear reinstatement, re-extinction and approach-avoidance task-AAT) one to three months later. For the cathodal group extinction was preceded by cathodal tDCS, whereas for the sham group, extinction was preceded by sham tDCS. During the 3 days, fear responses were measured by SCRs (except during tDCS stimulation and AAT), self-report ratings on valence, arousal, contingency and expectancy, and AAT (cf. Fig 1).

#### 2.3.1 Day 1

The habituation phase consisted of 8 non-reinforced presentations of the CSs. The acquisition phase consisted of a partial reinforcement procedure at 75% – i.e., 12 out of 16 presentations of the CS+ were paired with the auditory US. When the CS+ and US were paired, the presentation of the US overlapped with the last 2 seconds of the CS+. The CS− was never paired with the US. In habituation and acquisition, stimuli were presented for 4s over a white screen, followed by a jittered inter-stimulus interval (ISI; 10-12s), during the presentation of a black fixation cross over a white screen.

#### 2.3.2 Day 2

Participants that successfully acquired the fear response were randomly assigned to the cathodal or the sham group in a 2:1 ratio to compensate for cathodal stimulation variability.

In the extinction-learning phase, participants were asked to verbally recall the CS+. The session consisted of 16 CS+ and 16 CS− trials. Per trial, the fixation cross (ISI; 10-12s) was followed by the presentation of the CS+ or the CS− for 16s. The US was not presented during extinction.

#### 2.3.3 Day 3

Participants were invited to a follow-up session one to three months later. Sessions started by asking participants to verbally recall the color of the CS+. The reinstatement phase consisted of four consecutive unsignaled USs for 2s each (jittered ISI = 1-20s). The re-extinction phase started immediately after reinstatement and was similar to day 2 extinction session.

### Transcranial Direct Current Stimulation (tDCS)

The tDCS session was delivered offline and was 20min long. A constant current of 1mA^5^ and .04mA/mm^2^ current density was delivered through a tDCS 1-channel stimulator (TCT Research Limited, Hong Kong). We placed the tDCS cathode electrode over the rDLPFC (F4), according to the International 10-20 EEG System (45), and the anode electrode extra cephalically over the contralateral deltoid (22; see Fig S1 in Supplementary Materials). We used two electrodes of 24.75cm^2^ wrapped in saline soaked sponges (0.9% sodium chloride). Current was ramped up and ramped down during the first and last 30s of stimulation (43). At the end of the tDCS session, participants were instructed to report any adverse effects (cf. Table S1 in Supplementary Materials).

### Psychological questionnaires

Day 1 baseline psychological assessment was answered in a computer screen, using a keyboard, and included the following instruments: Anxiety Sensitivity Scale – 3-PT (46); Behavioral Symptoms Inventory (47); State Anxiety Inventory (STAI-1) and Trait Anxiety Inventory (STAI-2; 48). In day 2 and day 3, participants answered STAI-1 (48).

### Other self-report measures

We used Lang’s (49) Self-Assessment Manikin (SAM) scales to assess arousal and valence. Arousal ranged from one (*highly calm*) to nine (*highly excited*) and valence ranged from one (*highly unpleasant*) to nine (*highly pleasant*). We used a customized scale to assess contingency for the CSs/US association, from 0% (the CS was never paired with the US) to 100% (the CS was always paired with the US), in steps of 25%. To assess expectancy of the US presentation after each CS presentation, we used a customized scale ranging from zero (*I was sure the sound was not coming)*, to nine (*I was sure the sound was coming*).

The rating scales were presented on the screen using E-Prime and answered using the keyboard.

### Skin conductance responses

In day 1 and 3, Powerlab26T finger electrodes (MLT116F; ADInstruments, Ltd., Dunedin, New Zealand) were attached to the medial phalanges of the index and middle fingers of the left hand and connected to a galvanic skin response Amplifier (FE116; ADInstruments, Ltd., Dunedin, New Zealand). Data was collected at a rate of 5Hz, filtering out frequencies above 50Hz. The signal was pre-processed in MATLAB (2013, The MathWorks, Inc., Natick, Massachusetts, United States) using in-house scripts.

### Approach-avoidance task (AAT)

We used a symbolic task to assess implicit approach-avoidance components of fear (40). We presented a white frame either in a portrait or landscape position, either at the top or at the bottom of the screen, containing one of the CSs, for 2s. Participants used the keyboard to move the manikin as fast as possible, towards or away from the frame, according to its orientation and regardless the stimulus that was placed inside the frame. The manikin appeared for 1.5s, followed by the frame, and participants’ response (ISI=2s). Incorrect responses were followed by a red cross feedback (0.5s) and slow responses by an attention-call after 13s. Each participant completed two blocks – one where they were instructed to move the manikin towards the frame when it was in landscape orientation, and the other where participants were instructed to move the manikin away from the frame when it was in portrait orientation. The order of the blocks was counterbalanced across participants. The task included four practice trials per block using a grey square instead of colored squares. Each block had eight conditions, in a two (stimulus: CS+ vs. CS−) by two (frame orientation: portrait vs. landscape) by two (position of the frame: top vs. bottom) design and included a total of 40 experimental trials. We collected reaction times and accuracy (34). During the AAT, the headphones were not placed.

## Results

For the main analysis, we excluded 2 participants that did not successfully acquire a fear response according to contingency ratings. Other 3 participants were excluded due to missing data on day 1 contingency (caused by a recording error). A total of 43 participants completed the experiment (mean age = 20.42 ± 4.99; mean education years = 13.00 ± 1.90), of which 27 were randomly assigned to the tDCS group and 16 to the sham group. Groups did not differ in what concerns age, education and psychological questionnaires at baseline (cf. Table 1). To perform the analysis, we applied Greenhouse-Geisser correction when the sphericity assumption was not fulfilled according to Mauchly’s Test.

**Table 1.** Sample statistics at baseline. Mean values for baseline socio-demographic information and psychological assessment. No differences between groups were found. US: Unconditioned stimuli; ASI-3-PT: Anxiety Sensitivity Scale Portuguese version; BSI (GSI): Global index for symptoms intensity of the Behavioral Symptoms Inventory; STAI 1: State Anxiety Inventory; STAI 2: Trait Anxiety Inventory. Cathodal: tDCS cathodal stimulation group; sham: tDCS sham group. The standard errors of the mean (SEM) are within parenthesis.

|  | tDCS group<br>n = 27 | Sham group<br>n = 16 | *p* | *df* |
| --- | --- | --- | --- | --- |
| **Age, Years** | 19.81 (2.56) | 21.44 (7.53) | .416 | 17.08 |
| **Education, Years** | 13.00 (1.78) | 13.00 (2.13) | 1.00 | 41 |
| **US Intensity, %** | 93.34 (3.50) | 92.07 (4.20) | .290 | 41 |
| **ASI-3-PT** | 26.93 (13.09) | 21.25 (10.97) | .153 | 41 |
| **BSI (GSI)** | .03 (.01) | .03 (.01) | .492 | 40 |
| **STAI 1** | 2.34 (.32) | 2.32 (.19) | .771 | 41 |
| **STAI 2** | 2.42 (.20) | 2.44 (.28) | .815 | 40 |

Note. Mean values are depicted. The standard errors of the mean (SEM) are within parenthesis. US: Unconditioned stimuli; ASI-3-PT: Anxiety Sensitivity Scale Portuguese version; BSI (GSI): Global index for symptoms intensity of the Behavioral Symptoms Inventory; STAI 1: State Anxiety Inventory; STAI 2: Trait Anxiety Inventory.

### Self-Reports

In day 1, the results of the independent *t*-tests suggest comparable fear responses between groups (cathodal or sham) after habituation and acquisition for each stimulus (CS+ or CS−; t < 1). Two-way repeated measures ANOVA with stimulus (CS+ or CS−) as within-subjects factor, and group (cathodal or sham) as between-subject factor, show no interaction between stimuli and experimental group, both after habituation for reported arousal (*F* (1, 41) = .091, *p* = .764, *ηp*^*2*^ = .002) and valence (*F* (1, 33) =.005, *p* = .942, *ηp*^*2*^ = .000), and after fear acquisition for reported arousal (*F* (1, 41) = .279, *p*= .600, *ηp*^*2*^ = .007), valence (*F* (1,36) = 2,33, *p*= .136, *ηp*^*2*^ = .061), contingency (*F* (1, 41) = .00, *p* = .989, *ηp*^*2*^ = .000), and expectancy (*F* (1, 20) = .491, *p*=.492, *ηp*^*2*^ = .024). Note that due to registration errors, we lost the post-acquisition expectancy ratings from fifteen participants of the cathodal group and six participants of the sham group. Notwithstanding the importance of expectancy ratings, this measure was not used to set fear conditioning criteria, thus not critically influencing following procedures. After acquisition, a main effect of stimuli was statistically significant for arousal (*F* (1, 41) = 138.58, *p* < 0.001, *ηp*^*2*^ = .772), valence (*F* (1, 36) = 120.77, *p* < 0.001, *ηp*^*2*^ = .770, expectancy (*F* (1, 20) = 163.75, *p* < .001, *ηp*^*2*^ = .891), and contingency (*F* (1, 41) = 1240.89, *p* < 0.001, *ηp*^*2*^ = .968), such that the CS+ was rated as triggering increased arousal, having an increased negative affect, being more frequently paired with the US, and leading to increased expectancy of US presentations, when compared to the CS−, confirming fear acquisition across groups (cf. Fig S3-A in Supplementary Material).

In day 2, two-way repeated measures ANOVA, with stimulus (CS+, CS−) as within-subjects factor, and group (cathodal, sham) as between-subjects factor showed that after tDCS session but before extinction, there was no interaction between stimuli and experimental group for arousal (*F* (1, 41) = 1.97, *p* = .168, *ηp*^*2*^ = .046) and valence *F* (1, 41) = 3.13, *p* = .084, *ηp*^*2*^ = .071). However, the expected main effect of stimuli (due to day 1 fear acquisition) was present for arousal (*F* (1, 41) = 19.79, *p* < 0.001, *ηp*^*2*^ = .33) and valence (*F* (1, 41) = 29.97, *p* < 0.001, *ηp*^*2*^ = .42), with the CS+ still triggering increased arousal and leading to increased negative affect compared with the CS−. Accordingly, there is no evidence for immediate cathodal tDCS effect as participants scored the stimuli in a similarly way. After extinction, the interaction was still not statistically significant for arousal (*F* (1, 40) = .64, *p* = .427, *ηp*^*2*^ = .016), valence (*F* (1, 41) = 2.01, *p* = .164, *ηp*^*2*^ = .047), and expectancy ratings (*F* (1, 40) = .092, *p* = .763, *ηp*^*2*^ = .002). However, the main effect of stimuli with increased responses to the CS+ was still present and statistically significant for arousal (*F* (1, 40) = 11.35, *p* = 0.002, *ηp*^*2*^ = .22), valence (*F* (1, 41) = 9.70, *p* = 0.003, *ηp*^*2*^ = .19) and expectancy (*F* (1, 40) = 52.12, *p* < .001, *ηp*^*2*^ = .57). No main effect of group was found for arousal, valence, or expectancy. For contingency ratings, Wilcoxon Signed Rank Test, showed no main effect of stimuli (*Z* = .00; *p* = 1), and Mann Whitney test for group differences, showed that the mean rank differences between the CS+ and the CS− were also not statistically significant (*U* = 208.00, *p* =.441; cathodal group mean rank = 22.30; sham group mean rank = 21.50). These results show that participants still expected the CS+ to be paired with the aversive sound (meaning that the declarative associative memory CS+/US was intact despite extinction), along with a correct report of the absence of the aversive stimulus during extinction. Again, there is no evidence for short-term cathodal tDCS effect over extinction.

In day 3 (1 to 3 months after extinction), after recall but before re-extinction, two-way repeated measures ANOVA with stimulus (CS+, CS−) as within-subjects factor, and group (cathodal, sham) as between-subject factor showed results similar to day 2 post extinction, with no interaction between stimuli and experimental group for arousal (*F* (1, 41) = .645, *p* = .427, *ηp^2^* = .015), and valence (*F* (1, 41) = 1.046, *p* = .313, *ηp*^*2*^ = .03). Also, before re-extinction, a main effect of stimuli was present for arousal (*F* (1, 41) = 7.18, *p* = .011, *ηp^2^* =.149), and valence (*F* (1, 41) = 9.30, *p* = .004, *ηp*^*2*^ = .185), but no main effect of group for arousal (*F* (1, 41) = 1.02, *p* = .319, *ηp*^*2*^ =.024), and valence (*F* (1, 41) = .673, *p* = .417, *ηp*^*2*^ = .016). After re-extinction, there was still no statistically significant interaction between group and stimuli for arousal (*F* (1, 39) = .39, *p* = .536, *ηp*^*2*^ = .010), valence (*F* (1, 41) = .67, *p* = .420, *ηp*^*2*^ = .016), and expectancy (*F* (1, 41) = 1.71, *p* = .440, *ηp*^*2*^ = .015). The main effect of stimuli that was present before, disappeared after re-extinction for arousal (*F* (1, 39) = 1.20, *p* = .280, *ηp*^*2*^ = .030) and for valence (*F* (1, 41) = 2.26, *p* =.141, *ηp*^*2*^ = .052).

Regarding the expectancy ratings though, a main effect of stimuli was still present (*F* = (1, 41) = 27.35, *p* < .001, *ηp*^*2*^ = .400), indicating a pervasive expectancy of the US when presented with the CS+. No main effect of group was present for arousal (*F* (1, 39) =.019, *p* = .891, *ηp*^*2*^ = .000), valence (*F* (1, 41) = .14, *p* =.707, *ηp*^*2*^ = .003), or expectancy (*F* (1, 41) = .563, *p* = .458, *ηp*^*2*^ = .014). Mann-Whitney test for contingency reports, showed that the median rank differences between the CS+ and the CS− were not statistically significant between groups (*U* = 216.00, *p* = 1); and the Wilcoxon Signed Rank Test showed no effect of stimuli (*Z* = .00; *p* = 1) either. In sum, results indicate that self-reports across groups were equivalent, with a perseverance of declarative stimuli discrimination, despite the correct contingency learning (cf. Fig S4 in Supplementary materials).

### Skin conductance responses

In day 1, for habituation and acquisition phases, independent samples *t*-tests showed no differences between groups, per trial (cf. Fig S3-B in Supplementary material). Two-way repeated measures ANOVAs showed no interaction between groups (cathodal or sham) and the trial order of the SCRs differentials for habituation (*F* (6, 246) =.56, *p* = .762, *ηp*^*2*^ = .013), and for acquisition (*F* (8.993, 368,730) =.61, *p* = .787, *ηp*^*2*^ = .015). No main effects were found during habituation for trial order (*F*(6, 246) =.80, *p* = .570, *ηp*^*2*^ = .019) or for group (*F* (1, 41) =.28, *p* = .602, *ηp*^*2*^ = .007); nor during acquisition for trial order (*F* (8.993, 368,730) =1.29, *p* = .240, *ηp*^*2*^ = .031) or group (*F* (1, 41) =.016, *p* = .901, *ηp*^*2*^ = .000). In day 2, there was also no interaction between group and trial order (*F* (14, 574) = 1.36, *p* = .166, *ηp*^*2*^ = .032), and no main effects were found for stimuli (*F* (14, 574) =1.76, *p* = .041, *ηp*^*2*^ = .041), or for group (*F* (1, 41) =.87, *p* = .355, *ηp*^*2*^ = .021). Day 2 SCR results suggest that there were no short-term differences between groups along extinction, in what concerns the autonomic fear response.

In day 3 (1 to 3 months after extinction), we tested *extinction retention* using two-way repeated measures ANOVAs with stimuli (CS+ or CS−) as within subject factor, and group (cathodal or sham) as between subjects factor and found no statistically significant interaction (*F* (1, 41) = 2.57, p = .117, *ηp*^*2*^ = .059), no main effect of stimuli (*F* (1, 41) = 1.67, *p* =.203, *ηp*^*2*^ = .039), and a significant main effect of group (F (1, 41) = 5.24, *p* = .027, *ηp*^*2*^ = .113), with the cathodal group showing increased SCRs differentials. This result showed that tDCS may have a detrimental effect in long-term fear savings. However, when running a group by trial (2×15) repeated measures ANOVA across the re-extinction session, no interaction was found (*F* (9.586, 393.025) =1.0, *p* = .445, *ηp*^*2*^ = .024), and no main effects were present for the trial order (*F* (9.586, 393.025) =.89, *p* = .537, *ηp*^*2*^ = .021) or for group (*F* (1, 41) = .388, *p* = .537, *ηp*^*2*^ = .049), suggesting no evidence of cathodal tDCS impact on SCRs across the re-extinction process. Also, 1mA cathodal tDCS has a long-term detrimental effect in the fear memory savings towards de CS+ (however easily surpassed by the overall re-extinction results), which does not happen after sham tDCS.

### Approach-Avoidance Task

To explore differences in AAT performance across groups we performed a three-way repeated measures ANOVA with CS (CS+ and CS−) and response (approach and avoidance) as within-subjects factor and group (cathodal and sham) as between-subjects factor.

As expected, the three-way interaction was statistically significant (*F* (1, 41) = 5.97, *p* = .019, *ηp*^*2*^ =.127), meaning that there were group differences in AAT performance. To explore these differences, we followed up by two-way repeated measures ANOVAs using stimuli and response as within-subjects’ factors, for each group separately. We found a significant cross-over interaction for the sham group (*F* (1, 15) = 4.72, *p* = .046, *ηp^2^* = .239) but not for the cathodal group (*F* (1, 26) = 3.34, *p* = .079, *ηp^2^* =.114). That is, for the sham group, the effect of the stimuli differs with the type of response, although no main effects were found (differences between stimuli in avoidance RTs: *t* (15) = 1.39, *p* = .185; 95% CIs [−13.55, 64.43]; differences between stimuli in approach RTs: *t* (16) = −.76, *p* = .460; 95% CIs [−59.58, 28.33]). For the cathodal group there were no significant results for the two-way ANOVA. However, we further explored the responses in a similar manner as in the sham group. Follow-up planned contrasts for the cathodal group revealed a marginal difference between the avoidance RTs to the CS+ and the avoidance RTs to the CS− (*t* (26) = −2.00, *p* = .056; 95% CIs [−66.86, .97]). That is, in the cathodal group participants were overall faster to avoid the CS+ than to avoid the CS−. Because this difference did not survive corrections for multiple comparisons (*t* (26) = −2.00, *p* =.28; 95% CIs [−66.86, .97]) and because there was no statistically significant interaction, the results should be interpreted with caution. Furthermore, for the cathodal group we found a difference between the approach and the avoidance responses to the CS− (*t*(26) = −4.86, *p* < .0001, 95% CIs [−118.96, −47.89], Bonferroni corrected) with overall faster approach RTs, suggesting a positive valence attributed to the CS− (two-tailed; Fig 2A).

**Fig 2.**
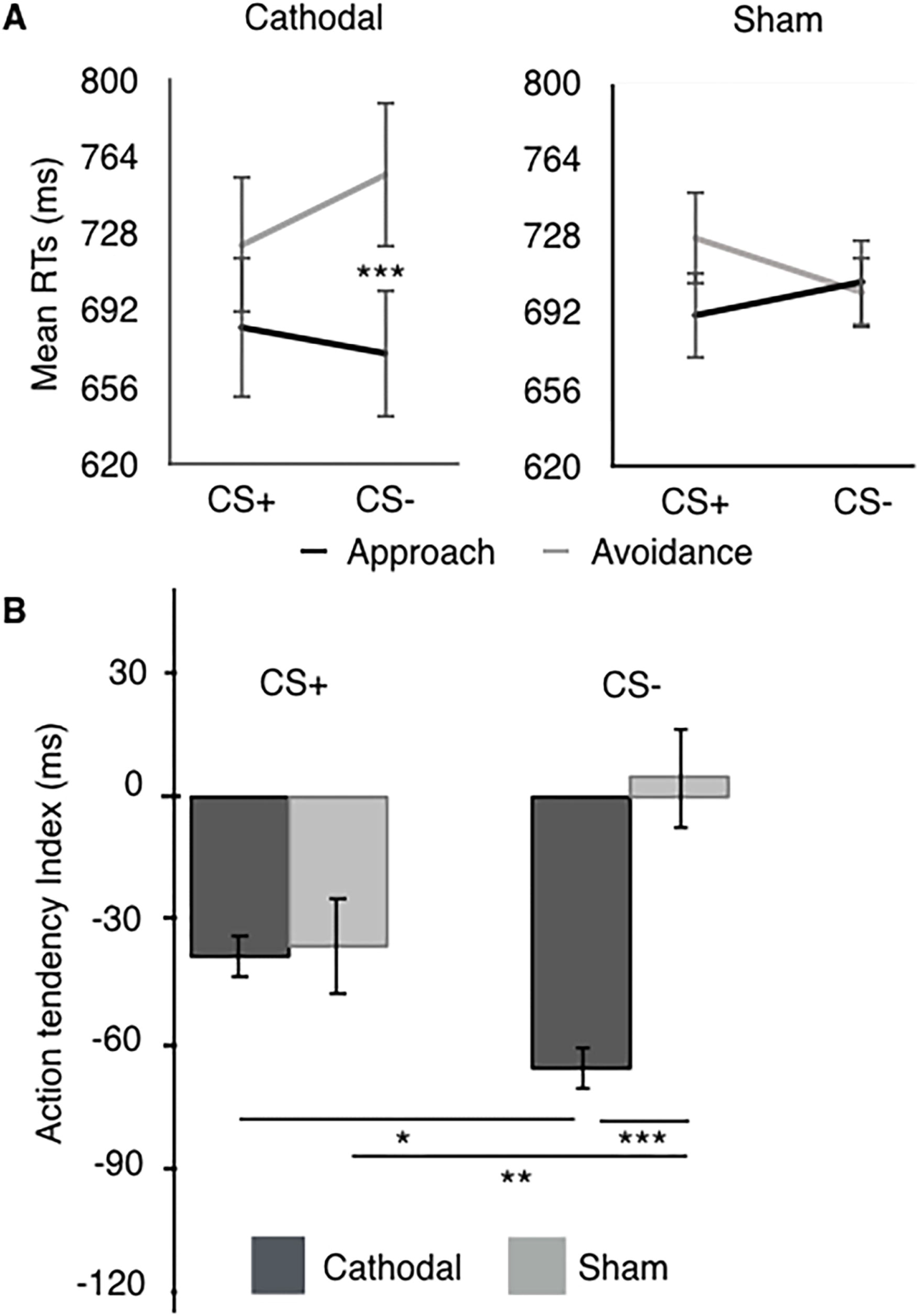
Approach-avoidance task (AAT) results. A) Approach and avoidance responses per experimental group. *** p < .001; CS+: conditioned stimuli; CS−: non-reinforced or control stimuli; cathodal: tDCS cathodal stimulation group; sham: tDCS sham group. Error bars represent standard errors of the mean (SEM). (Single column image, grey scales)

Following previous literature (40), we computed the avoidance tendency index (the time to approach subtracted to the time to avoid) for each stimulus, in each group. Two-way repeated measures ANOVA, with the stimuli (CS+, CS−) as within-subjects factor and the group (cathodal, sham) as between-subjects factor showed an interaction between stimuli and experimental group with a moderate effect size (*F* (1, 41) = 5.97, *p* = .019, *ηp^2^* = .127). Univariate analyses suggest that whereas there was no difference between groups regarding the avoidance tendency towards the CS+ (*F* (1, 41) = .006, *p* = .937, *ηp^2^* = .000), there was a difference between groups in the avoidance tendency towards the CS− (*F* (1, 41) = 12.04, *p* = .001, *ηp^2^* = .227). In fact, a moderate effect size is present for the estimated difference between groups, with the avoidance tendency presented by participants of the cathodal group towards the CS− following the trend of positive stimuli (positive bias), whereas for the sham group it remains equal to zero (cf. Fig 2B and Table S3 in supplementary materials). Although caution is required for the interpretation of the results in the absence of an interaction between stimuli and response, this result suggests that whereas tDCS may have no long-term effect over the avoidance tendency towards the conditioned stimuli, it may have an effect over the avoidance tendency towards the neutral stimuli, leading to a disambiguation of its value.

## Discussion

In this study, we assessed whether cathodal tDCS (assumed to induce neural inhibition) over the rDLPFC enhances the efficacy of fear extinction procedures across three distinctive components of the fear response – the autonomic response (SCRs), the subjective experience (self-reports) and the implicit avoidance tendencies (AAT index).

In a sample of 34 women, we found no evidence of cathodal tDCS short-term impact in the way participants perceive threatening cues, according to its autonomic component. Surprisingly, offline cathodal tDCS shows a long-term negative effect on SCRs, as illustrated by our extinction retention test, wherein participants in the cathodal group showed increased fear memory savings to the conditioned stimulus. Abend et al. (20), within a similar procedure aimed at disrupting reconsolidation and using low-frequency oscillations alternating current (AC) to produce long-term depression effect (LTD), found a similar result. Like AC, cathodal tDCS is expected to decrease neural activity by depolarizing apical dendrites and hyperpolarizing the somatic regions of the pyramidal cells under the cathode electrode (50). Together, these results show no evidence of effect for cathodal tDCS over short-term fear conditioned SCRs, and evidence of a detrimental effect of cathodal tDCS over long-term fear and SCRs.

However, both cathodal tDCS and AC, resulted in the preservation of adequate stimuli discrimination, whereas anodal stimulation leads to a fear generalization effect to the CS− (18,20). In generalization, the conditioned fear response spreads to similar perceptual stimuli. Subsequently, same class stimuli become aversive and individuals learn that they need to be avoided. In our study, we confirm previous literature findings concerning the use of AC (20) and show that fear generalization does not occur after cathodal stimulation, strengthening the hypothesis that generalization is a polarity adverse outcome exclusive of anodal stimulation.

Furthermore, our study shows how different fear measures may suggest distinctive conclusions. Whereas affective, contingency, expectancy and self-reported state anxiety show no effect, SCRs suggest increased long-term fear memory savings for the CS+, after cathodal stimulation. This asynchrony was previously linked to the lack of overlap between the mechanisms behind each measure. In fact, the expectancy of harm or the intolerance to uncertainty that emerges during the extinction procedure by the absence of the US are differently translated by explicit ratings and SCRs (51,52).

Finally, our results on avoidance tendencies support SCRs data, showing that the way cathodal tDCS benefits extinction, may occur not by enhancing the fear response to the threatening cue, but through the mitigation of generalization to similar stimuli. This means that although cathodal stimulation has no direct effect over the new learning about the CS+, it may have an effect over the CS−. We found was that the tendency to avoid the CS+ has similarly decreased after classical extinction with or without cathodal tDCS. However, for the cathodal group there seems to be a positive bias towards the CS−, which is not present in the sham group. On the contrary, for the sham group, the avoidance tendency to the neutral cue increased beyond expected. According to the AAT index, it seems then that tDCS may have indeed an effect in neutralizing the fearful response pattern to both threat cues and neutral cues, limiting generalization.

A generalization effect of the fear response to the CS− was previously found as a result of anodal tDCS (18,20) and our study suggests that this effect may be polarity-dependent. If cathodal stimulation is effective in reducing anxiety symptoms in case reports (22,23), it would be expected to be effective in boosting its analogous experimental extinction procedures. Indeed, although according to the most common fear measures there is no effect of cathodal stimulation on extinction, there is an effect of cathodal stimulation on the stimuli discrimination process associated with implicit avoidance tendencies. Accordingly, ridding off generalization to perceptively similar stimuli can be the mechanism by which cathodal tDCS succeeds to mitigate anxiety related symptoms. As generalization is a known transdiagnostic factor across anxiety disorders, and one of the hardest features to overcome in therapy, it may be that cathodal tDCS has a beneficial potential as an add-on enhancer to extinction (which is in contrast with the disadvantageous generalization effect that comes along with anodal tDCS).

Here we show that cathodal stimulation over the rDLPFC enhanced the safety learning about similar neutral cues (the CS−). According to previous literature, it may be so by directly decreasing the DLPFC activity, known to respond to stimuli similar to the CS+ (53), or by indirectly enhancing the vmPFC response to the CS− during extinction (54). Regardless the actual mechanism of action, our study suggests that cathodal tDCS may be altering simultaneous activity of a whole circuit, interfering with the balance between excitation and inhibition of different regions within the fear network, boosting discriminative processes (35).

A single cathodal tDCS session may indeed impact both short- and long-term extinction (1 to 3 months) and the fear system components. According to the literature cathodal tDCS of 1mA can lead to both immediate and prolonged polarity-specific effects (8). Whereas the immediate effects of tDCS are due to the modulation of the membrane resting potential (by affecting ion channels and altering its homeostasis), the after-effects rely on the modulation of enduring synaptic plasticity, interfering with long-term potentiation (LTP) and long-term depression (LTD). These are activity dependent plasticity mechanisms that result in persistently enhanced or reduced synaptic transmission. Furthermore, the effect of tDCS over LTP and LTD is triggered by prolonged stimulation durations (such as a single 20min session) and is also moderated by concurrent task-specific synaptic plasticity (14). That is, the tDCS effect is task-dependent in that its effect is moderated by concurrent behavioral interventions such as associative learning (of which fear conditioning is one example). Particularly, as already shown in animal models, the combination of cathodal tDCS with behavioral fear conditioning modifies the strength of the associative learning by changing the functional properties of the network (55).

Furthermore, we know that tDCS modulates long-distance connectivity to subcortical structures (11). Hence, previous fMRI studies show that the DLPFC downregulates fear responses through projections to the vmPFC that in turn inhibit the amygdala during extinction learning. However, how does tDCS affect the amygdala is still not clear. Previous literature showed increased vmPFC activity during classical extinction (17) but decreased when extinction occurs within the reconsolidation window (25). Whereas in our study we found that cathodal tDCS putatively occurring during the reconsolidation window, may reduce the fear index/avoidance tendency in the long-term, Mungee et al. (21) found no effect for cathodal tDCS in SCRs. Whilst in our study we asked participants to verbally recall the CS+ followed by classical extinction (days 2 and 3), in Mungee et al. (21) there was no information update because participants did not go through a complete extinction procedure following tDCS. However, according to seminal studies, this new information is a necessary condition for the update of the CS+/US association (24).

On the other hand, the atypical reminder we used seemed to be not effective in triggering reconsolidation, and the expected update of the associative learning concerning the CS+ was not achieved (see the increased fear recovery in the beginning of day 3, indexed by SCRs). Whether cathodal tDCS has successfully modulated reconsolidation boundaries concerning the conditioned fear response towards the CS−, by decreasing the vmPFC participation during extinction, while simultaneously enhancing the discriminative learning, is not straightforward in light of the reconsolidation theory. In this sense, our results may further contribute to the recent debate around reconsolidation theory robustness and consistency when applied to fear conditioning procedures (e.g., 56,57).

Results seem to suggest that after cathodal tDCS, discrimination between threatening and neutral cues is enhanced and a positive bias towards potentially ambivalent stimuli is present in the implicit behavioral fear system. This transference of positive affect is illustrated not only by the absence of avoidance tendencies towards the fear-conditioned stimulus (CS+), but also by a positive bias towards the neutral stimulus (CS−) in day 3, only present for the cathodal group. In fact, the sham group showed that beyond the absence of avoidance tendencies towards any of the stimuli, there was a positive bias towards the CS− after extinction.

We used tDCS to boost extinction and to persistently eliminate conditioned fear responses. Although tDCS does not bring any particular improvement to declarative memory associated measures (self-reports and SCRs), the generalization of the implicit behavioral fear response to perceptively similar stimuli seems to be decreased. As such, adding cathodal tDCS may enhance the efficacy of extinction-based treatments by inhibiting the generalization effect – a therapeutic benchmark of anxiety disorders such as generalized anxiety disorder and phobias. We believe thus, that the use of cathodal tDCS as an add-on strategy may be a promising strategy, although further confirmatory studies should be conducted in the future.

## Acknowledgements

We thank Soares, M.J and Gerardo, B. for data collection and preprocessing.

## Funding

AGA is supported by the Foundation for Science and Technology, Portugal and Programa COMPETE [grants numbers SFRH/BD/80945/2011, PTDC/MHC-PAP/5618/2014 (POCI-01-0145-FEDER-016836); http://www.poci-compete2020.pt/]. JA is supported by the Foundation for Science and Technology, Portugal and Programa COMPETE [grants numbers PTDC/MHC-PAP/5618/2014 (POCI-01-0145-FEDER-016836), PTDC/MHC-PCN/3575/2012, PTDC/MHC-PCN/0522/2014, PTDC/MHC-PCN/0522/2014, PTDC/MHC-PCN/6805/2014; https://www.fct.pt/index.phtml.en]. The Cognitive and Behavioral Center for Research and Intervention of the Faculty of Psychology and Educational Sciences of the University of Coimbra is supported by the Portuguese Foundation for Science and Technology and the Portuguese Ministry of Education and Science through national funds and co-financed by FEDER through COMPETE2020 under the PT2020 Partnership Agreement [UID/PSI/01662/2013; https://www.portugal2020.pt/Portal2020]. The Psychology Research Centre of the University of Minho, is supported by the Portuguese Foundation for Science and Technology and the Portuguese Ministry of Education and Science through national funds and co-financed by FEDER through COMPETE2020 under the PT2020 Partnership Agreement (POCI-01-0145-FEDER-007653). The Proaction Laboratory and PTDC/MHC-PAP/5618/2014 (POCI-01-0145-FEDER-016836) supported this research.

## Data Availability

The datasets generated during and/or analyzed during the current study are available at https://osf.io/rmnq3/?view_only=1a9110a2a178453789df74c885a995b6.

## Author contributions

AGA, OFG, PB, AMK, MKA designed the study. AGA and RG collected data and analyzed the data. AGA wrote the main manuscript; All authors discussed and reviewed the manuscript.

## Competing interests

The authors have declared that no competing interests exist.

