## Supporting information for "The effect of cathodal tDCS on fear extinction: a cross-measures study"

This document includes:

- Supplementary Text

Methods

Participants

Stimuli

Pre-processing data

Analytic strategy

Supplementary Figures: S1, S2

Results

Psychological questionnaires

- Supplementary Tables: S1, S2, S3

- Supplementary Figures: S1, S2, S3, S4, S5

- References

### Methods

**Participants**

In this study, We tested only female participants due to known gender differences concerning electrodermal activity ^1^, and fear conditioning responses^2^.

**Stimuli**

The intensity of the US was individually set in the first session, using a dummy aversive sound (item 276 from the International Affective Digitized Sound System)^3^ and a Visual Analog Scale for pain^4^ to measure discomfort level during sound test (minimum level = 90 dB; maximum level = 96 dB). The US presentations were delivered using noise canceling headphones, and participants were instructed to report the sound intensity level that led to maximum discomfort without pain.

### Data collection and pre-processing data

***Skin conductance responses.*** To protect for device drifts we performed pre- baseline correction using the mean value of the 0.5s immediately before stimuli onset^5^. Because data was not normally distributed, we z-transformed SCR values^6^. We adopted the last observation carried forward imputation method for missing values for no responses and artifacts, as this is the most conservative option. We kept only participants for which the total number of missing values was less than half of the total trials per stimulus. SCRs analyses were ran over computed CSs differentials between CS+ and CS- per trial. The CS- was always set to be the first stimulus to be present in each session and it was disregarded, as it was assumed to be an orientation response^1^. To even the number of trials for each CSs we disregarded the last CS+ within each session. We calculated SCRs on a trial-by-trial basis using data from the first 3.5s after the onset of each stimulus.

Approach-avoidance task (AAT). In this task, RTs are thought to be dependent on the relationship between two independent types of responses: the explicit task-related response (i.e., to approach or avoid the frame according to its orientation), and the implicit fear-related response (i.e., to approach the neutral stimulus and to avoid the feared stimulus). If the action tendency component of the fear response is not fully eliminated (after extinction and re-extinction training procedures), there should be an effect of the implicit fear-related responses on the task-related approach/avoid responses. Specifically, RTs should be faster when these two responses are congruent (e.g., participants have to avoid the frame because of its orientation, and the frame contains the CS+), than when these two responses are incongruent (e.g. participants have to approach the frame because of its orientation, and the frame contains the CS+)^7^. From the RTs we calculated the fear index, subtracting the RT for the fear-related response of avoidance from the fear-related response of approach for each CS. A negative fear index mirrors the absence of avoidance tendencies, such that participants are faster to approach than to avoid, whereas a positive fear index shows the reverse tendency.

We excluded incorrect trials (6.83%) and trials with response times bellow 200ms and above 3000ms (4.30%) and used median RTs to compute fear indexes per stimulus ^8^. For an example of one trial of the AAT, see Fig S1.

### Analytic strategy

### **Psychological questionnaires.** To discard baseline differences that could otherwise influence results, we used independent t-tests to compare groups concerning each psychological measure. Further, independent samples t-test were used to observe tDCS impact over state anxiety, comparing group scores before extinction (day 2) and reinstatement (day 3).

***Self-reports.*** To make sure of the equivalency of groups at baseline we ran independent sample t-tests for the affective ratings of arousal and valence after habituation, and on the affective, contingency and expectancy ratings after acquisition. To understand the effects of stimulus and experimental group over self-report ratings, we performed a two-way repeated measures ANOVA with stimulus (CS+, CS-) as within-subjects factor, and group (cathodal, sham) as between-subject factor for day 1 (post-acquisition), day 2 (pre- and post-extinction) and day 3 (pre- and post-re-extinction).

***Skin conductance responses.*** In order to confirm similarity between groups during habituation and acquisition phases, we ran independent samples t-tests per trial. To better understand fear, as indexed by the SCRs during acquisition (day 1), extinction (day 2), and re-extinction (day 3), we performed two-way repeated measures ANOVAs for the CSs differentials, with trial (1 to 15) as within-subjects factor, and group (cathodal, sham) as between-subject factor. To observe fear recovery after reinstatement in day 3, we computed an independent samples t-test for the first responses to the CS+.

Approach-avoidance task (AAT). Three-way repeated measures ANOVA followed by simple contrasts allowed us to understand the interaction between stimulus (CS+ or CS-) and fear related response (approach or avoidance) as within subject factors, and tDCS group (cathodal or sham) as the between subject factor. In case of no interactions we further explore the data using simple contrasts to better understand within and between group patterns. Following previous literature ^8^, we further ran a two-way mixed ANOVA for the fear index followed by univariate analysis.

**Results**

### As expected from previous literature, participants reported few mild adverse effects to tDCS stimulation, mostly after cathodal stimulation. Nonetheless no differences between groups were found according to Mann-Whitney U test for independent samples (Table S1).

**Psychological questionnaires**

### In day 1, independent samples t-tests for each psychological measure showed no baseline differences between groups (*t* < 1). Similarly, state anxiety (STAI 1) was not different between groups in day 2 and 3 (*t* < 1), suggesting that tDCS stimulation did not impact self-reported symptoms.

**Self-Reports**

Due to registration errors we lost the post-acquisition expectancy ratings from fifteen participants of the cathodal group and six participants of the sham group. Final analysis included the available data sets. Results in day 1 suggest comparable fear responses between groups after habituation and acquisition for each stimulus (*t* < 1). As expected, we found no interaction between stimuli and experimental group regarding affective ratings after the habituation (arousal: *F* (1, 41) = .091, *p* = .761; valence: *F* (1, 33) = .005, *p* = .942). After fear acquisition, we found no interaction between stimuli and experimental groups for the level of reported arousal (*F* (1, 41) = .279, *p* = .600) but there was a main effect of stimuli due to fear acquisition, with participants reporting the CS+ as more arousing than the CS- (*p* < .001). Similarly, we found no interaction between stimuli and groups for the reported valence (*F* (1, 36) = 2,33, *p* = .136), but there was a main effect of stimuli, with participants reporting the CS+ as more unpleasant than the CS- (*p* < .001). For contingency and expectancy ratings we found no interaction (*F* (1, 41) = 0, *p* = .989; *F* (1, 20) = .491, *p* = .492, respectively), but a main effect of stimuli was present in both (*p* < .001) with participants reporting increased contingency ratings to the CS+/US association than to the CS-/US, and increased expectancy for the US when presented with the CS+ (for further details see Fig S1-A).

In day 2, after tDCS session and before extinction training, results showed the same trend as in day 1, with no interaction between stimuli and experimental group for arousal (*F* (1, 41) = 1,97, *p* = .168) and valence *F* (1, 41) = 3,13, *p* = .84), but a main effect of stimuli (*p* < .001) with increased arousal and unpleasantness reported to the CS+. After extinction, the interaction was still not statistically significant for arousal (*F* (1, 40) = ,64, *p* = .427), and valence (*F* (1, 41) = 2.01, *p* =.164) and there was a main effect of stimuli still present (arousal: *p* = .002; valence: *p* = .003). For the contingency ratings, there was no interaction between stimuli and experimental group (*p* = 1) and the main effect of stimuli disappeared, as expected after successful extinction training. Finally, no interaction was found for expectancy ratings (*F* (1, 40) = .092, *p* = .763), but a main effect of stimuli was still present (*p* < .001) showing that participants had an increased expectation of the US presentation when presented with the CS+, even after a long extinction procedure, and irrespectively of the tDCS group they were assigned to.

In day 3, before re-extinction the results showed no interaction between stimuli and experimental group for arousal (*F* (1, 41) = .645, *p* = .427), and valence (*F* (1, 41) = 1.046, *p* = .313), and the main effect of stimuli was present (arousal: *p* = .011; valence: *p* = .004), with participants rating the CS+ as eliciting increased arousal and unpleasantness. However, after re-extinction, besides no statistically significant interaction between stimuli and experimental group for arousal (*F* (1, 39) = .389, *p*= .536) and valence (*F* (1, 41) = .665, *p* = .420), the main effect of stimuli disappeared (*p* > .05), suggesting no effect of tDCS in that the CS+ and the CS- were affectively similar, in both groups. As happened in day 2, contingency ratings were equivalent across stimuli and groups (*p* = 1), and no main effect of stimuli was found. Nonetheless, the presence of a fear component can not be ruled out, as a main effect of stimuli was still present after re-extinction (*p* = .001), despite no interaction between stimuli and group for expectancy ratings (*F* (1, 41) =.61, *p* = .440), suggesting that participants preserved an increased expectancy of the US when presented with the CS+, irrespectively of the group they were assigned to (for detailed self-reports for the three days see Table S2).

**Skin conductance response**

### To observe the relation between the SCRs and the contingency accuracies at day 1, we conducted a contingency table analysis between the SCRs and the contingency ratings. We dichotomized the SCRs according to its common acquisition criteria (0.01µS was used as the cut-off value for discrimination vs. non-discrimination) and dichotomized the contingency variable into ‘high’ and ‘low’ accuracy (‘high’ was defined as the combinations of responses to the CS+/US > 50% [responses of 75% or 100%] with responses to the CS-/US = 0% [responses of 0%]; ‘low’ was defined as the combinations of responses to the CS+/US < 75s% [responses of 0% or 25% or 50%] with responses to the CS-/US ≥ 25% [responses of 25%, 50%, 75% or 100%). Fisher's exact test yields a p = .323 (*Ø =*.183, *p =* .230), meaning that there is no evidence that the two variables are associated. Of note, the contingency ratings of our participants do not illustrate the full spectrum of possible responses as they are already selected according to the acquisition/non-acquisition criteria at study entrance (CS+/US ≥ 50% and CS-/US < 50%)^9^.

### The literature has shown how different measures may represent different fear learning dimensions, and although SCRs are usually associated with declarative ratings of stimuli discrimination this is not always true, particularly when the rate of CS+/US pairing is intermediate, as happens in our study (75%)^10^. Also, contingency ratings are considered a requisite when establishing a subjective measure of successful acquisition of the fear response^9^. Together, these arguments led us to set the contingency ratings as the primary outcome to establish fear acquisition.

**Supplementary Tables**

| **Table S1**. *Reported tDCS stimulation adverse effects.* | | | |
| --- | --- | --- | --- |
|  | **cathodal** | **sham** | ***p*** |
|  | n = 27 | n = 16 |  |
| **Headache** | 3 | 0 | .172 |
| **Neck pain** | 0 | 0 | 1 |
| **Scalp discomfort** | 0 | 0 | 1 |
| **tingling** | 8 | 2 | .204 |
| **itching** | 5 | 0 | .070 |
| **bruning sensation** | 1 | 1 | .705 |
| **local skin redness** | 5 | 0 | .070 |
| **somnolence** | 7 | 2 | .301 |
| **attention deficit** | 1 | 0 | .441 |
| **mood changes** | 0 | 1 | .194 |
| **Others** | 1 | 0 | .441 |
| Note. Frequencies and significance *p* value from Mann-Whitney U for independent samples are depicted. cathodal: tDCS cathodal stimulation group; sham: tDCS sham group. | | | |

| **Table S2**. *Self-report ratings.* | | |  | |  |  | |  |  |  | |
| --- | --- | --- | --- | --- | --- | --- | --- | --- | --- | --- | --- |
|  | **Phase** |  | **CS+** | | | |  | **CS-** | | |  |
|  |  |  | **cathodal (n=27)** | **sham (n=16)** | | | ***p*** | **cathodal (n=27)** | | **sham (n=16)** | ***p*** |
| **Day 1** | habituation | arousal | 3.48 (.40) | 3.63 (.44) | | | .818 | 4.11 (.42) | | 4.06 (.49) | .942 |
|  |  | valence | 5.79 (.28) | 5.14 (.25) | | | .123 | 5.41 (.34) | | 5.15 (.34) | .643 |
|  | post-acquisition | arousal | 6.85 (.30) | 6.50 (.39) | | | .478 | 3.30 (.27) | | 3.25 (.43) | .924 |
|  |  | valence | 2.25 (.28) | 2.93 (.34) | | | .143 | 6.48 (.29) | | 6.31 (.41) | .734 |
|  |  | contingency | 3.93 (.07) | 3.81 (.10) | | | .364 | 1.11 (08) | | 1 (.00) | .185 |
|  |  | expectancy | 8.17 (.21) | 7.10 (.53) | | | .084 | 1.58 (.56) | | 1.20 (.42) | .600 |
| **Day 2** | pre-extinction | arousal | 4.85 (.37) | 5.81 (.28) | | | .072 | 3.78 (.34) | | 3.75 (.38) | .957 |
|  |  | valence | 4.37 (.36) | 3.25 (.34) | | | .**042** | 5.78 (.26) | | 6.00 (.37) | .618 |
|  | post-extinction | arousal | 4.33 (.33) | 4.81 (.38) | | | .360 | 3.81 (.34) | | 3.75 (.41) | .916 |
|  |  | valence | 5.26 (.27) | 4.69 (.31) | | | .181 | 5.70 (.30) | | 5.88 (.38) | .724 |
|  |  | contingency | 1.08 (.08) | 1.00 (.00) | | | .440 | 1.08 (.08) | | 1.00 (.00) | .431 |
|  |  | expectancy | 4.42 (.53) | 4.69 (.62) | | | .753 | 1.50 (.30) | | 2.00 (.55) | .389 |
| **Day 3** | pre-reinstatement | arousal | 4.63 (.34) | 5.38 (.38) | | | .164 | 3.89 (.41) | | 4.00 (.44) | .861 |
|  |  | valence | 4.63 (.34) | 4.50 (.38) | | | .808 | 5.41 (.30) | | 6.06 (.34) | .153 |
|  | post-extinction | arousal | 4.07 (.34) | 4.38 (.34) | | | .167 | 4.16 (.34) | | 3.94 (.41) | .559 |
|  |  | valence | 5.48 (.29) | 5.13 (.27) | | | .559 | 5.67 (.31) | | 5.75 (.31) | .692 |
|  |  | contingency | 1.00 (.00) | 1.00 (.00) | | | .413 | 1.00 (.00) | | 1.00 (.00) | .860 |
|  |  | expectancy | 2.63 (54) | 3.31 (.45) | | | .338 | .96 (.30) | | 1.06 (.30) | .829 |

Note. Mean values for arousal, valence, contingency and expectancy in day 1, 2 and 3. US: Unconditioned stimuli; CS+: conditioned stimuli; CS-: non-reinforced or control stimuli; cathodal: tDCS cathodal stimulation group; sham: tDCS sham group. In parenthesis are the standard errors of the mean (SEM).

| **Table S3.** *Approach and avoidance responses* | | | | | | |
| --- | --- | --- | --- | --- | --- | --- |
|  | **cathodal (n=27)** | |  | **sham (n= 16)** | |  |
|  | **CS+** | **CS-** | ***p*** | **CS+** | **CS-** | ***p*** |
| Approach | 684.00 (32.77) | 671,93 (30.11) | .787 | 690,72 (26.03) | 706.34 (26.50) | .667 |
| Avoidance | 722,41 (32.26) | 755,35 (34.53) | .056 | 726.81 (28.47) | 701.38 (22.39) | .488 |
| AAT Index | - 38.41 | -83.43 | 0.079 | -36.09 | 4.97 | **.005** |
| Note. Mean values for approach and avoidance responses, and for the AAT index (approach minus avoidance responses) are depicted. Differences between stimuli were found for the AAT index in the sham group. No differences between groups were found. CS+: conditioned stimuli; CS-: non-reinforced or control stimuli; Within parenthesis are the standard errors of the mean (SEM). Cathodal: tDCS cathodal stimulation group; sham: tDCS sham group; CS+: conditioned stimuli; CS-: control stimuli. | | | | | | |

**Supplementary Figures**

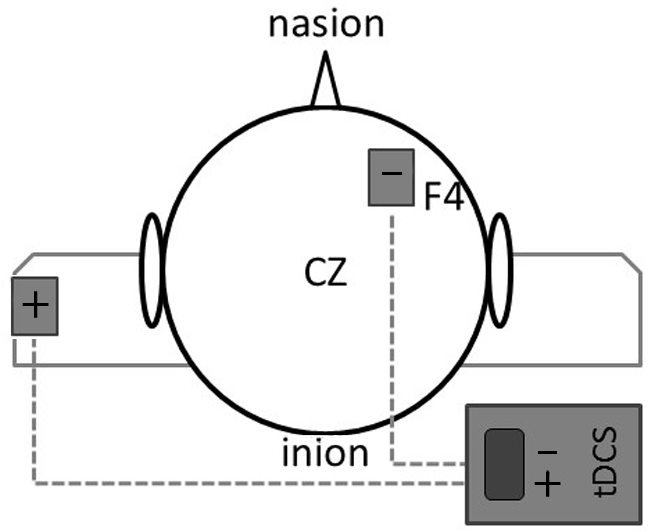

**Fig S1. tDCS montage.** Transcranial Direct Current Stimulation setting. The cathode electrode (represented by a minus sign) was positioned over F4 (i.e., right dorsolateral prefrontal cortex). The anode electrode (represented by a plus sign) was positioned over the left deltoid muscle; cz = vertex.

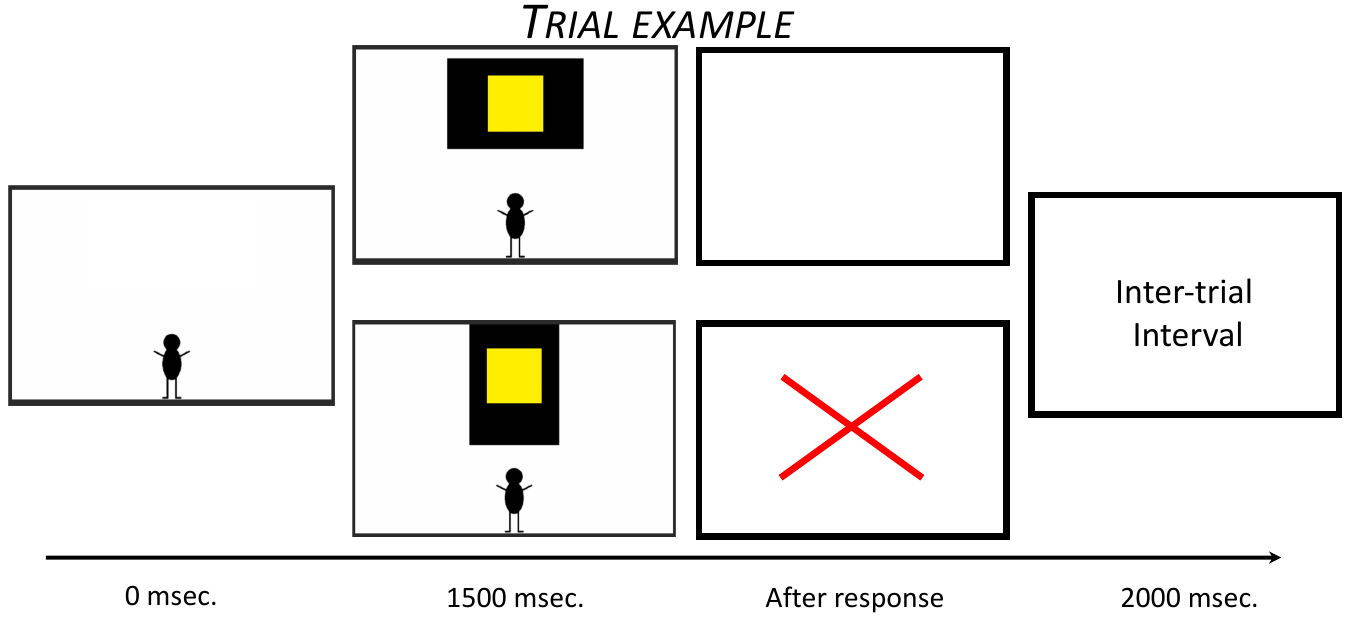

**Fig S2. AAT trial example.** Time-line for a trial example of the AAT.CS+: conditioned stimuli; CS-: non-reinforced or control stimuli

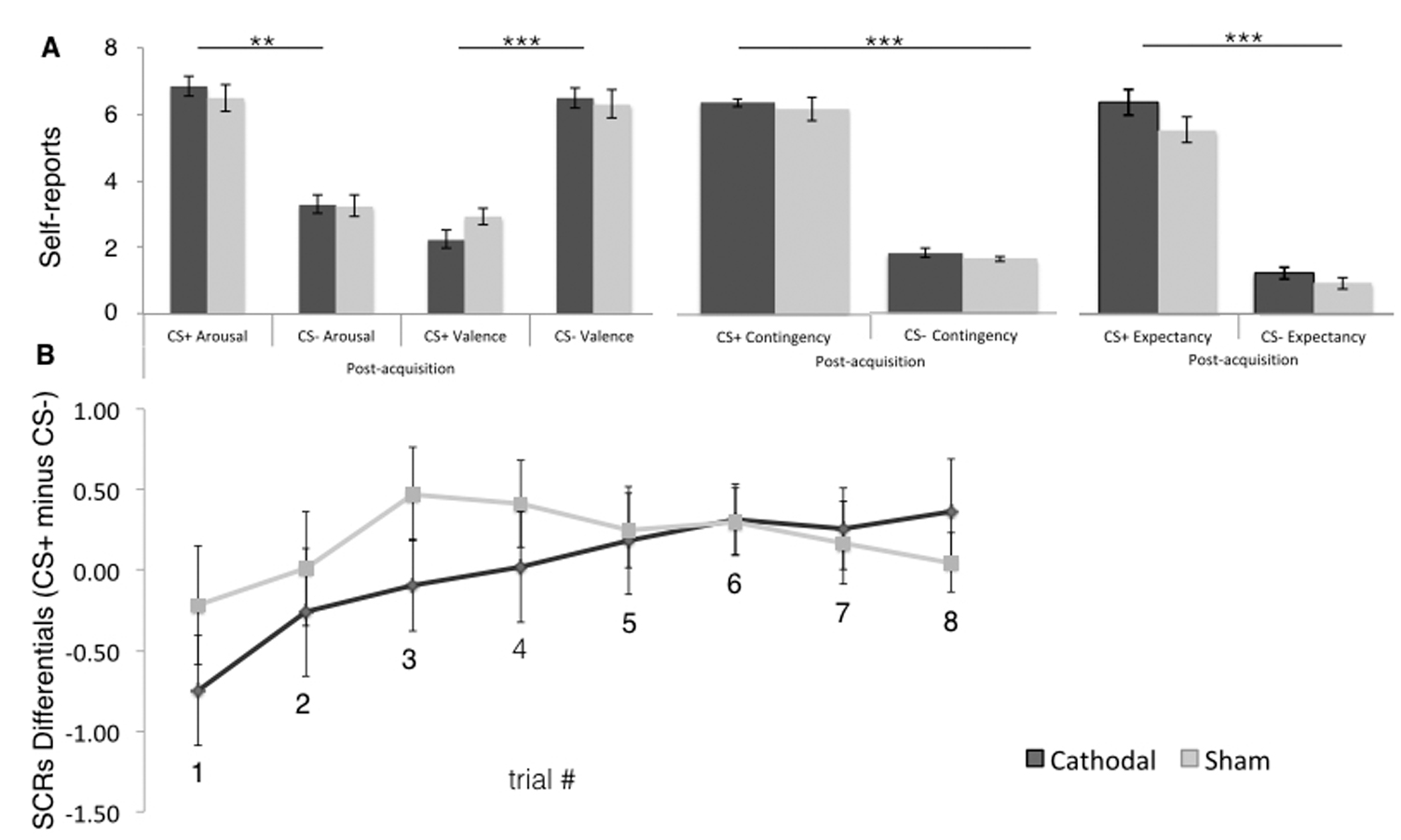

**Fig S3. Day 1 Self-reported fear acquisition measures.** Fear measures of A) affective self-reports of arousal and valence; contingency and expectancy for the CS+ and the CS-; B) skin conductance responses differentials, calculated by subtracting the CS- from the CS+ per trial. CS+: conditioned stimuli; CS-: non-reinforced or control stimuli; cathodal: tDCS cathodal stimulation group; sham: tDCS sham group. Error bars represent standard errors of the mean (SEM).

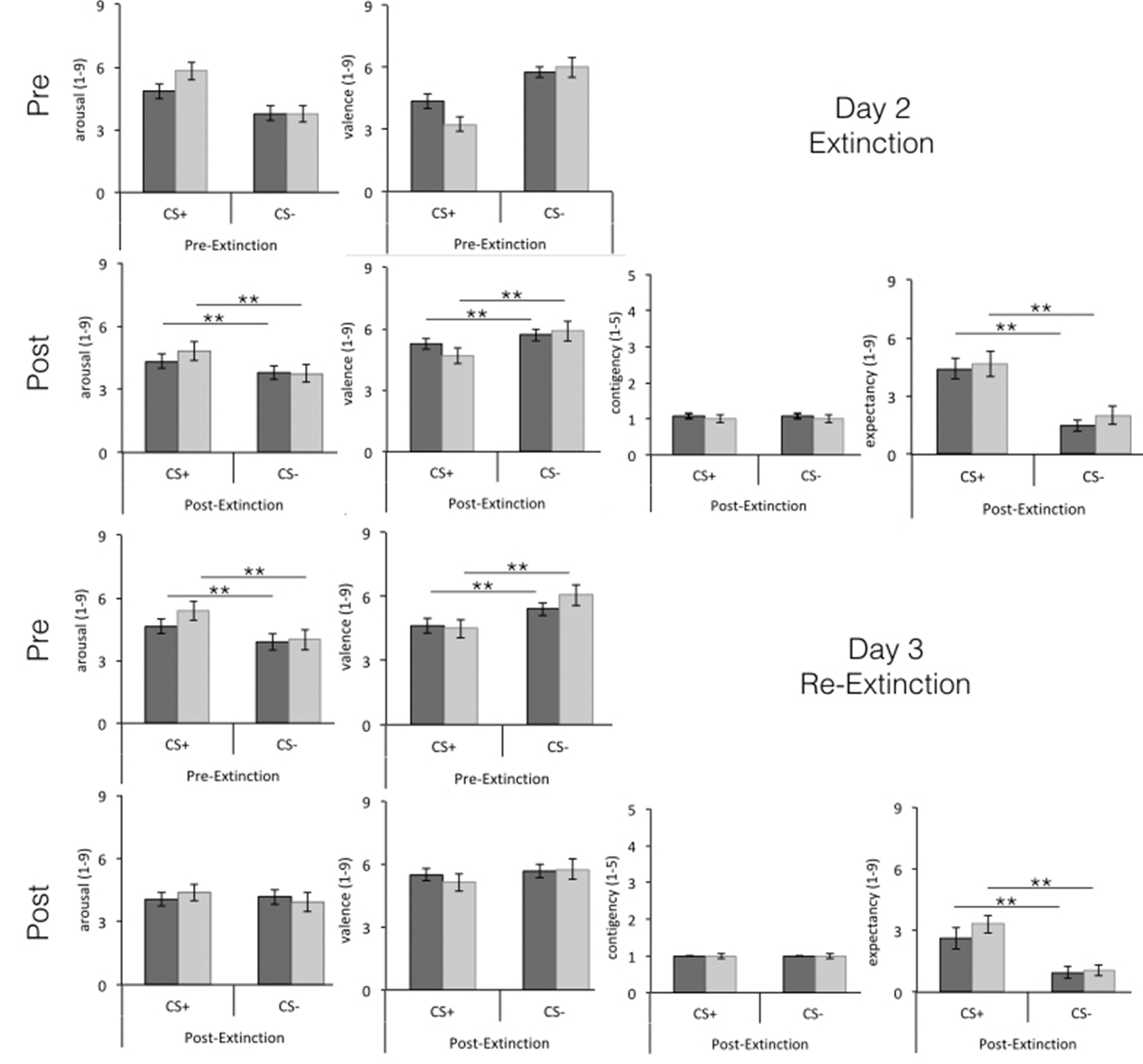

**Fig S4. Day 2 and Day 3 Self-reported fear acquisition measures.** Fear measures of affective self-reports of arousal and valence; contingency and expectancy for the CS+ and the CS-. CS+: conditioned stimuli; CS-: non-reinforced or control stimuli; cathodal: tDCS cathodal stimulation group; sham: tDCS sham group. Error bars represent standard errors of the mean (SEM).

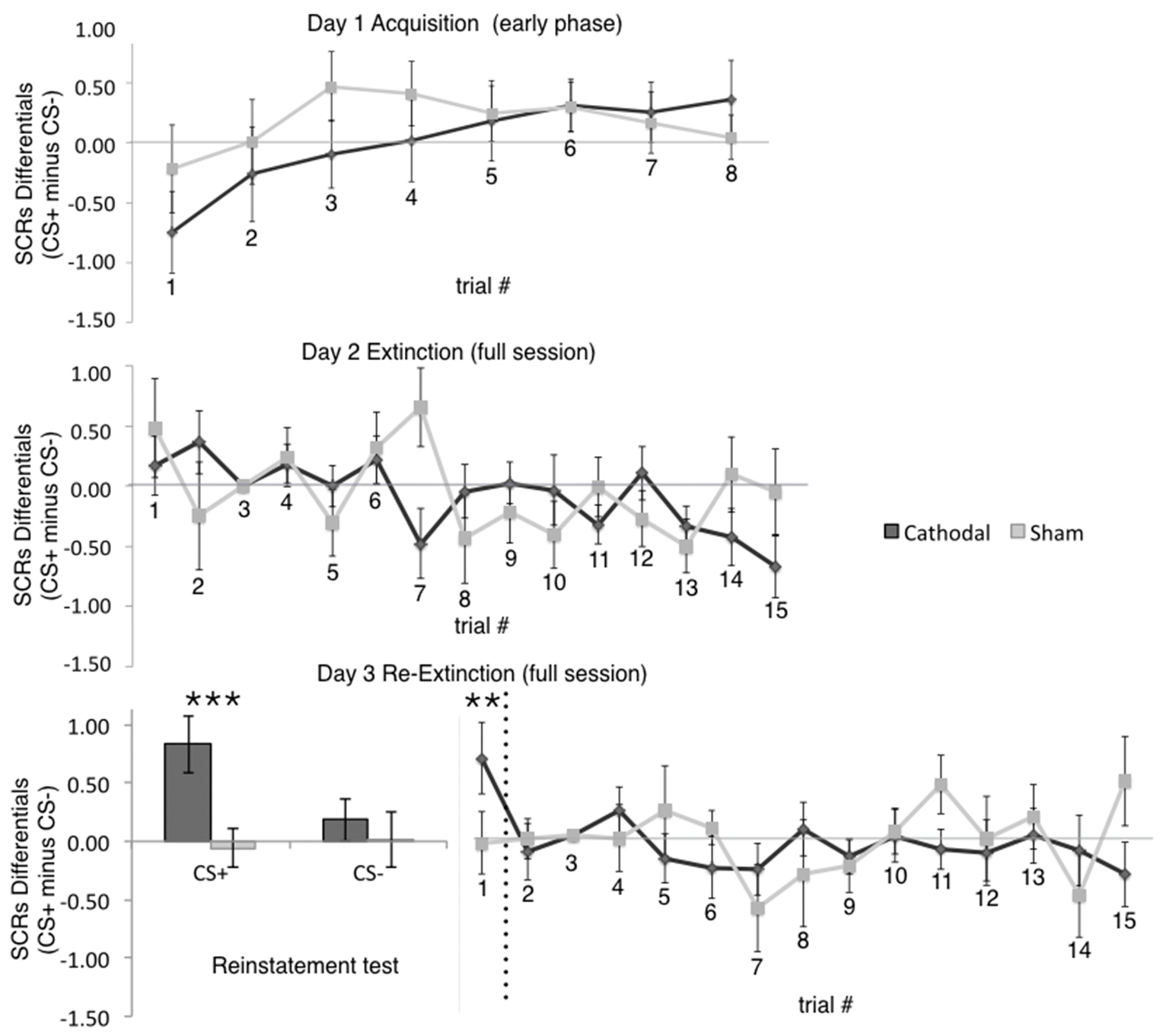

**Fig S5. Day 1, 2 and 3 SCRs differentials.** Skin conductance responses differentials (CS+ minus CS-) in day 1 (acquisition early phase), day 2 (extinction full session) and day 3 (reinstatement and re-extinction – full session). For better visualization, the histogram depicts the reinstatement test separately for the CS+ and for the CS-. CS+: conditioned stimuli; CS-: non-reinforced or control stimuli; cathodal: tDCS cathodal stimulation group; sham: tDCS sham group. Error bars represent standard errors of the mean (SEM).
