## Supplementary figures and images for "The effect of cathodal tDCS on fear extinction: a cross-measures study"

### FigureS1

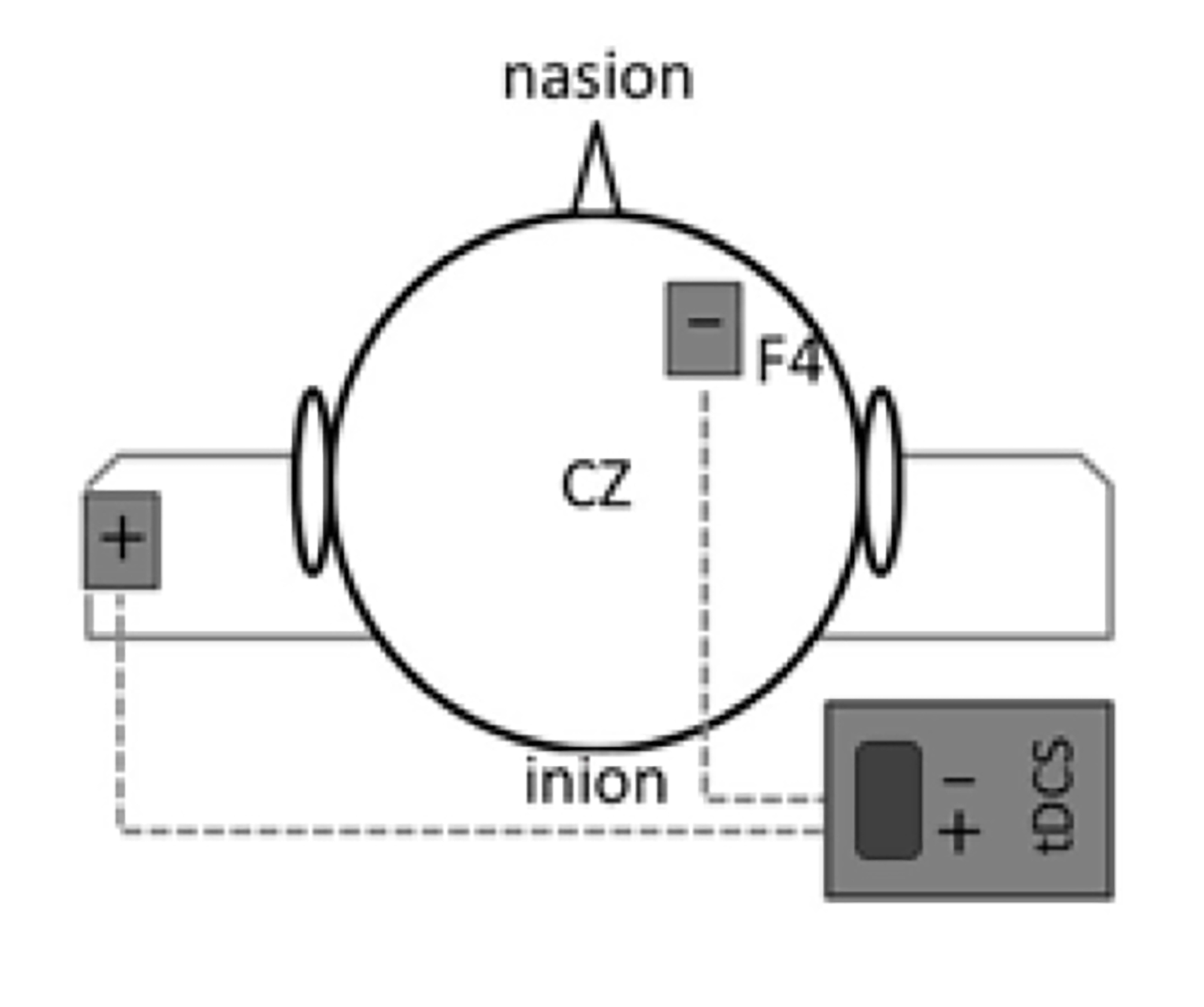

### FigureS2

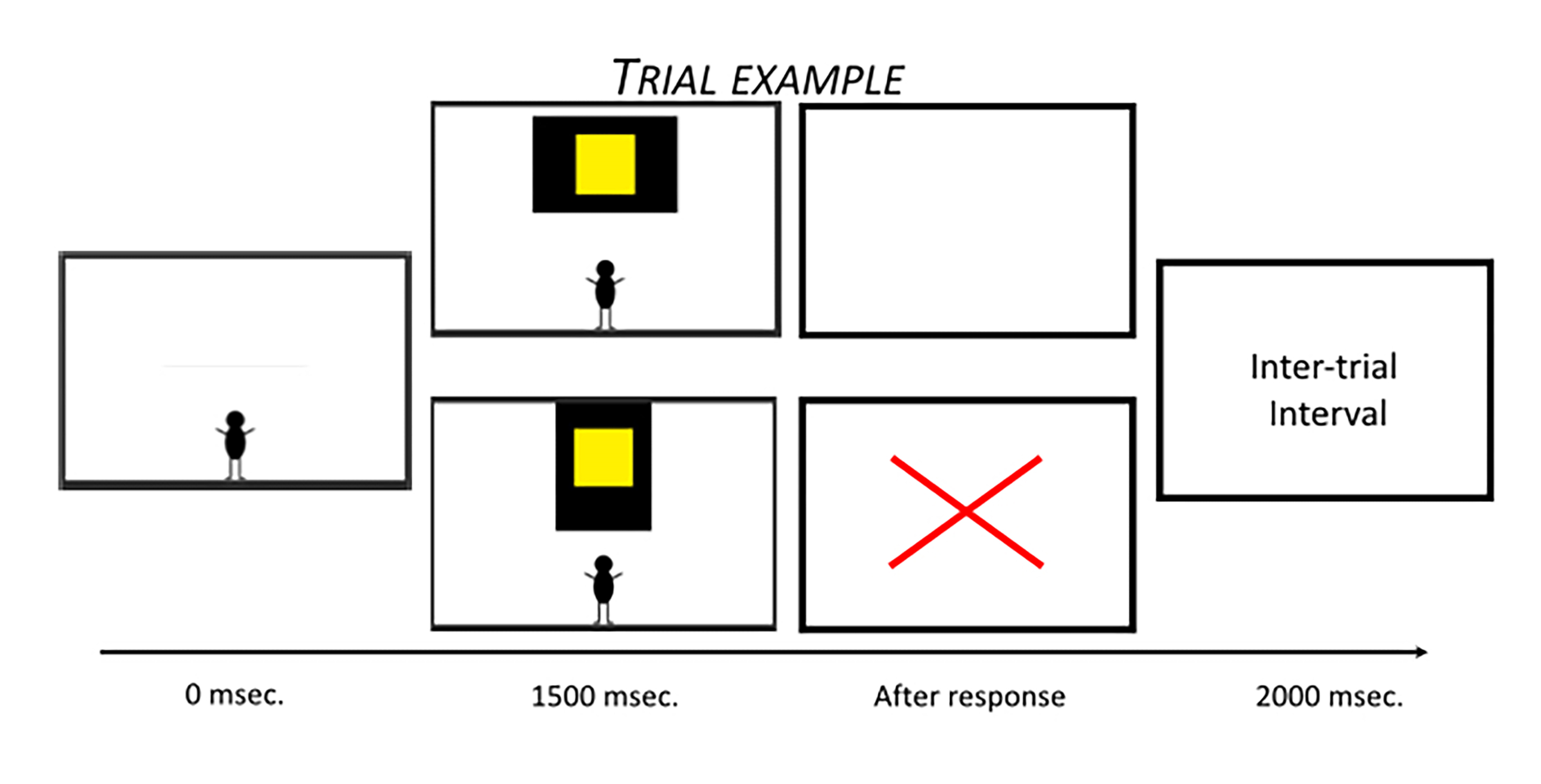

### FigureS3

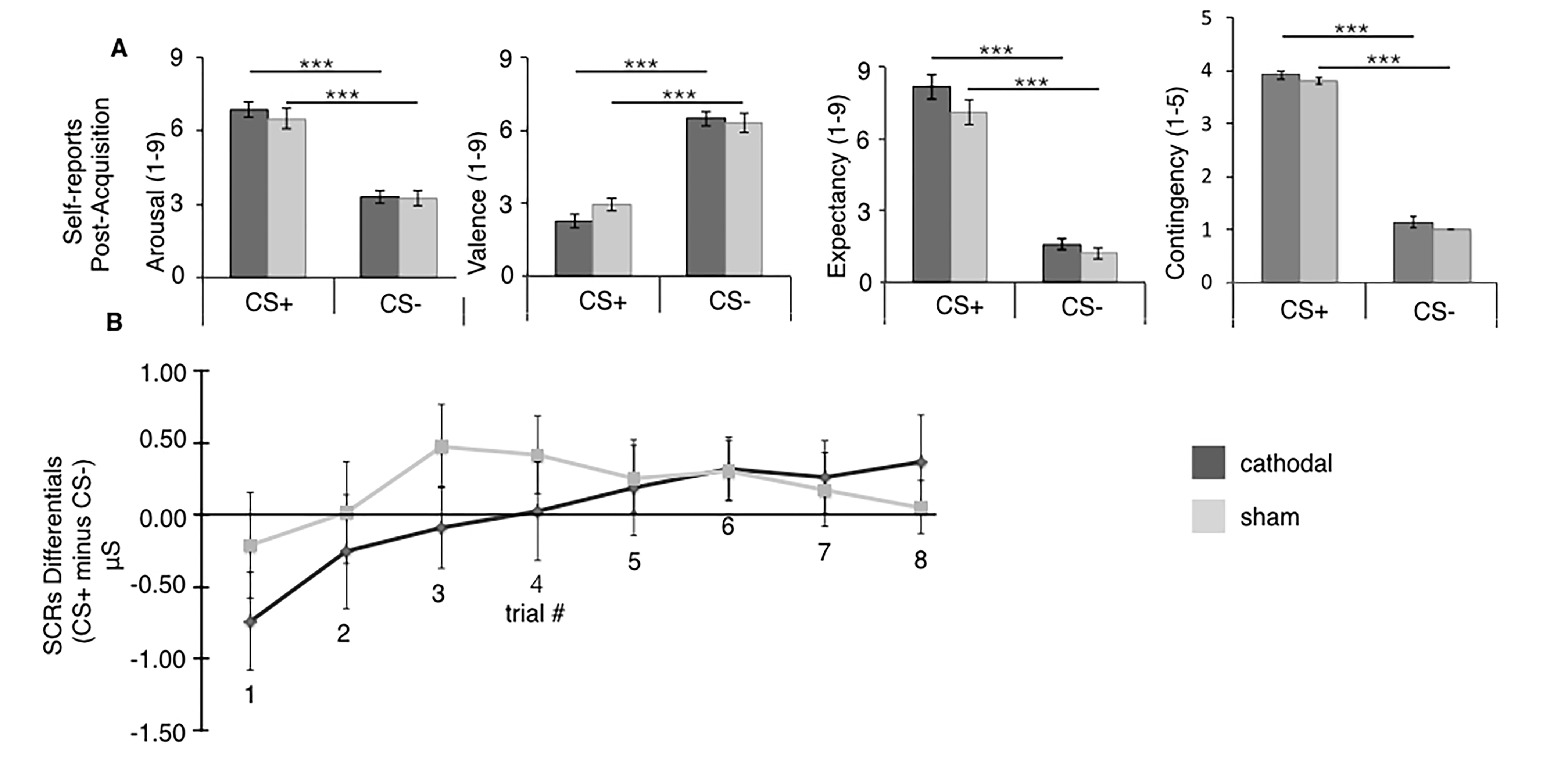

### FigureS4

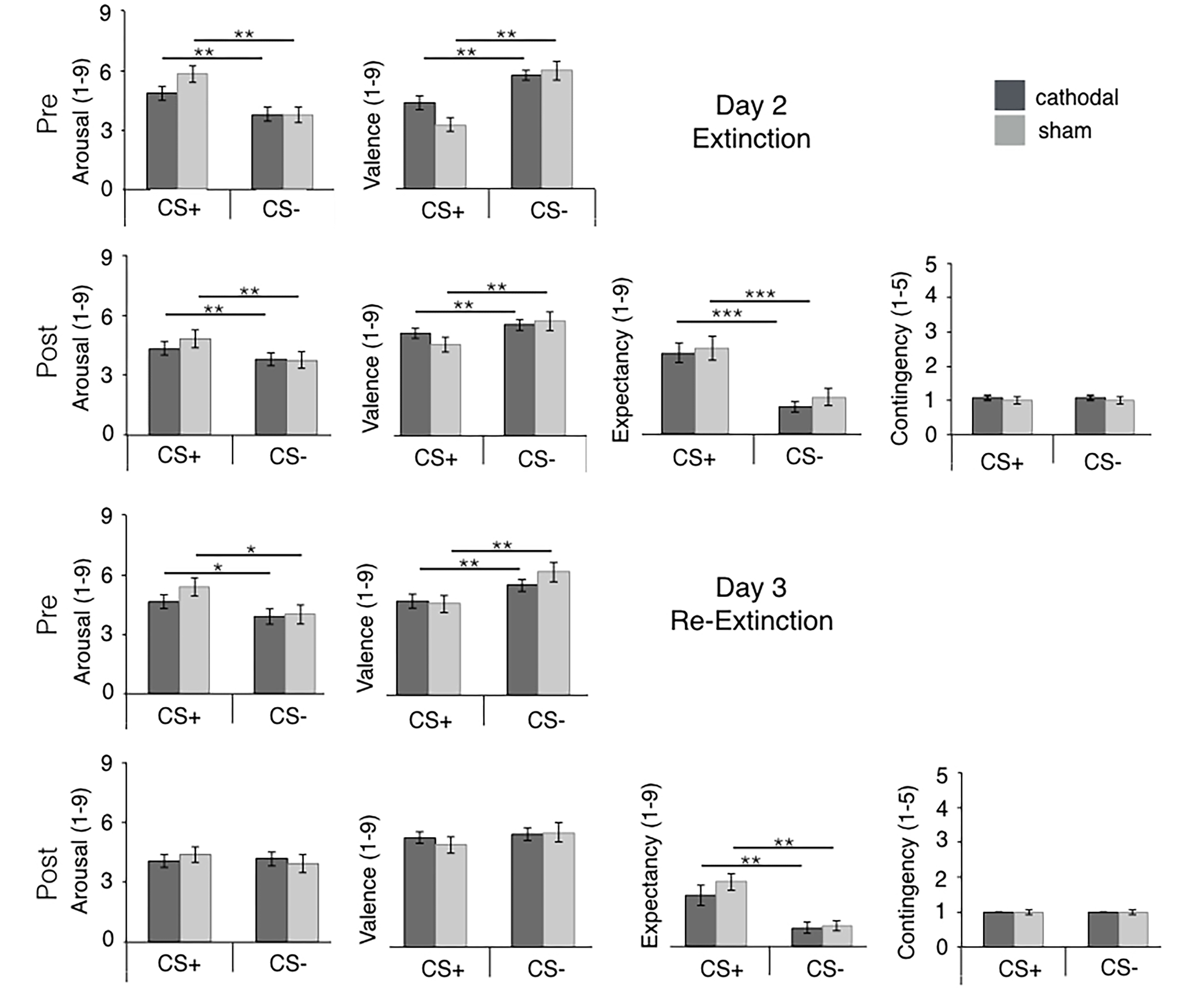

### FigureS5

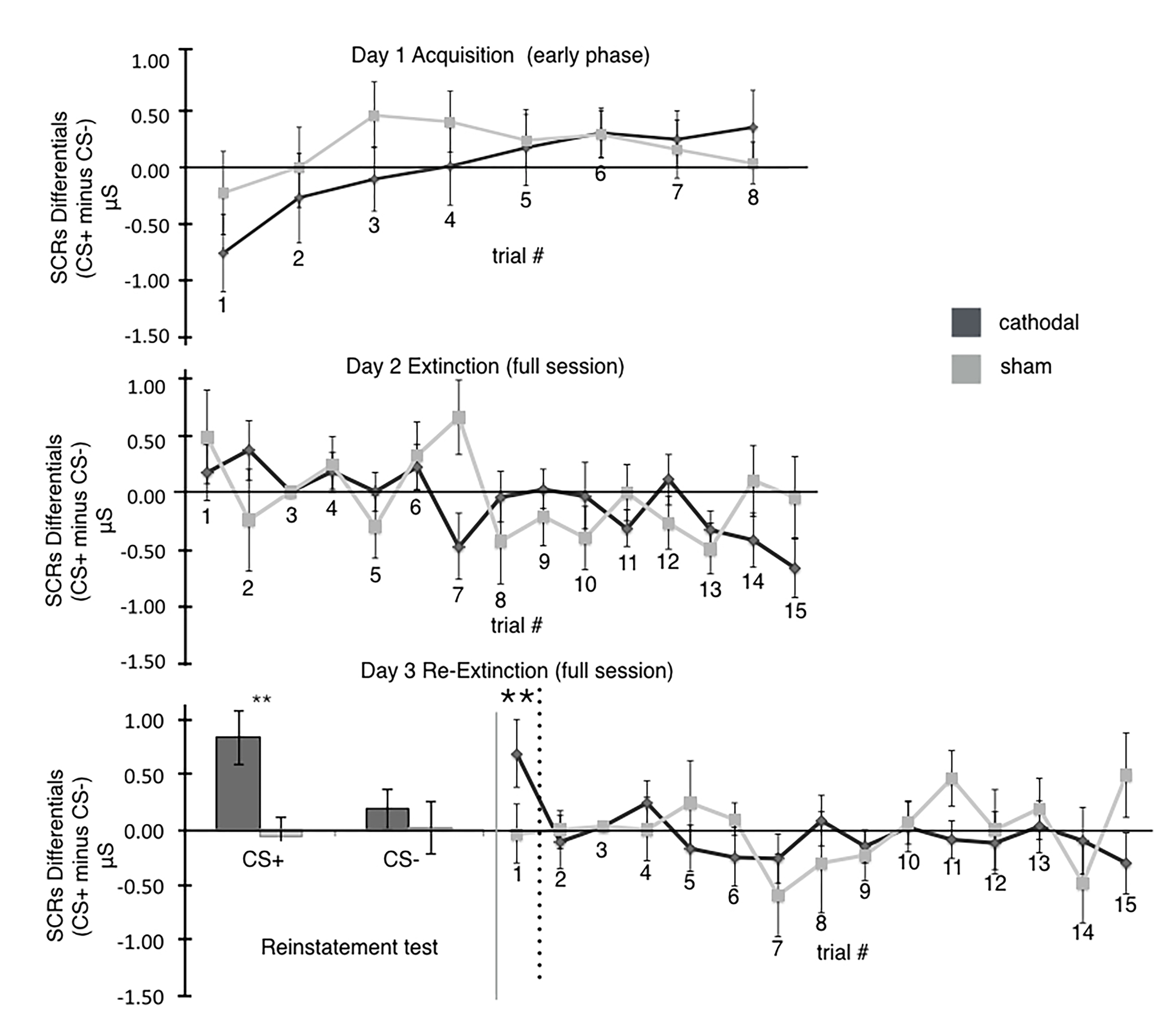
